# A population of neurons that produce hugin and express the *diuretic hormone 44 receptor* gene projects to the corpora allata in *Drosophila melanogaster*

**DOI:** 10.1101/2021.03.15.435215

**Authors:** Yosuke Mizuno, Eisuke Imura, Yoshitomo Kurogi, Yuko Shimadaigu-Niwa, Shu Kondo, Hiromu Tanimoto, Sebastian Hückesfeld, Michael J. Pankratz, Ryusuke Niwa

## Abstract

The corpora allata (CA) are essential endocrine organs that biosynthesize and secrete the sesquiterpenoid hormone, namely juvenile hormone (JH), to regulate a wide variety of developmental and physiological events in insects. Previous studies had demonstrated that the CA are directly innervated with neurons in many insect species, implying the innervations to be important for regulating JH biosynthesis in response to internal physiology and external environments. While this is also true for the model organism, *Drosophila melanogaster*, which neurotransmitters are produced in the CA-projecting neurons are yet to be clarified. In this study on *D. melanogaster*, we aimed to demonstrate that a subset of neurons producing the neuropeptide hugin, the invertebrate counterpart of the vertebrate neuromedin U, directly projects to the adult CA. A synaptic vesicle marker in the hugin neurons was observed at their axon termini located on the CA, which were immunolabeled with a newly-generated antibody to the JH biosynthesis enzyme JH acid *O*-methyltransferase (JHAMT). We also found the CA-projecting hugin neurons to likely express a gene encoding the specific receptor for diuretic hormone 44 (Dh44). Moreover, our data suggested that the CA-projecting hugin neurons have synaptic connections with the upstream neurons producing Dh44. To the best of our knowledge, this is the first study to identify a specific neurotransmitter of the CA-projecting neurons in *D. melanogaster*, and to anatomically characterize a neuronal pathway of the CA-projecting neurons and their upstream neurons.

## Introduction

An endocrine organ is responsible for biosynthesizing specific hormones to control animal development and physiology. In general, hormone biosynthesis in the endocrine organ is influenced by conditions of internal physiology and external environments, and regulates organismal homeostasis. In vertebrates, most endocrine organs are innervated with sympathetic and parasympathetic nerves, and the innervations play indispensable roles in the regulation of hormone biosynthesis (Schmidt-Nielsen, 1997).

Insect endocrine organs are also regulated by neurons that directly innervate the organs. One of the best examples is the prothoracic gland (PG), which biosynthesizes and secretes the principal insect steroid hormone ecdysone (Niwa and Niwa, 2014a; Pan et al., 2020), while the neurons produce prothoracicotropic hormone (PTTH) in the fruit fly *Drosophila melanogaster* (Kannangara et al., 2021; Malita and Rewitz, 2021; McBrayer et al., 2007; Niwa and Niwa, 2014b). Two pairs of cell bodies of the PTTH neurons are located in the brain hemispheres, extending their axons to the PG cells. PTTH is released at the synapses between PTTH neurons and the PG cells, stimulating the biosynthesis and secretion of ecdysone to regulate the timing of larval-to-pupal transition as well as body growth (McBrayer et al., 2007; Shimell et al., 2018). In addition to PTTH neurons, a subpopulation of serotonergic neurons is known to directly innervate the PG to regulate ecdysone biosynthesis in *D. melanogaster*. Cell bodies of the PG-projecting serotonergic neurons, called SE0_PG_ neurons, are located at the subesophageal zone (SEZ) and extend their long axons to the PG. The SE0_PG_ neurons are essential for controlling developmental timing in response to nutritional availability (Shimada-Niwa and Niwa, 2014). Recently, a subset of neurons producing the neuropeptide corazonin (Crz) has been reported to project to the PG in *D. melanogaster*, although the role of Crz neurons in ecdysone biosynthesis is still unclear (Imura et al., 2020).

Another endocrine organ in insects, the corpus allatum (plural: corpora allata (CA)), is also innervated with certain neurons. The CA are crucial organs for the biosynthesis and secretion of the sesquiterpenoid hormone named juvenile hormone (JH), which plays essential roles in a wide variety of aspects of insect development and physiology, including the larval-to-adult transition, oogenesis, epithelial morphogenesis, neuronal development, diapause, behavior, and longevity (Martín et al., 2021; Riddiford, 2020). Previous studies had suggested that neuronal inputs to the CA are essential for regulating JH biosynthesis and/or secretion, thereby regulating the biological events. For example, in the acridid grasshopper *Gomphocerus rufus*, and the locusts *Locusta migratoria* and *Schistocerca gregaria*, the CA are innervated with two morphologically distinct types of brain neurons (Virant-Doberlet et al., 1994). In the cockroach *Diploptera punctata*, nervous connections between the CA and brain are required for mitosis of the CA cells (Chiang et al., 1995). In the northern blowfly *Protophormia terraenovae* and the brown-winged green bug *Plautia stali*, neurons projecting to the CA are essential for inducing reproductive dormancy in short-day and low-temperature conditions (Matsumoto et al., 2013; Shiga et al., 2003). In case of *P. stali*, the CA-projecting neurons produce the neuropeptide *Plautia stali* myoinhibitory protein (Plast-MIP), which is considered to directly suppress JH biosynthesis in the CA (Hasegawa et al., 2020; Matsumoto et al., 2017). In *D. melanogaster*, at least two types of neurons had been reported to directly innervate the CA (Siegmund and Korge, 2001). These neurons, which seem to highly express a gene encoding the cell adhesion molecule Fasculin2, are involved in the epithelial movement of male genitalia by inhibiting JH biosynthesis during the pupal stage (Ádám et al., 2003). All the accumulated evidence together indicate direct neuronal innervation of the CA to be crucial for regulating JH biosynthesis in the latter. However, in almost all cases, except Plast-MIP, the neurotransmitter produced in the CA-projecting neurons has not been clarified. To further understand the neuroendocrine mechanism of JH biosynthesis in the CA, clarification and characterization of the neurotransmitters would be crucial.

Using *D. melanogaster*, a previous pioneering study had demonstrated that a subset of neurons producing the neuropeptide hugin (Hug), the invertebrate counterpart of vertebrate neuromedin U (Meng et al., 2002), extends the axons to the complex formed by the CA and corpora cardiaca (CC) in the adult insect (Melcher and Pankratz, 2005). However, which endocrine organ (the CA or CC) the Hug neurons project to still remains unclear. In this study, we aimed to provide an anatomical evidence of the subset of Hug neurons that directly project into the CA of *D. melanogaster*. The CA-projecting Hug neurons express a gene encoding the specific receptor for diuretic hormone 44 (Dh44), namely *Dh44-R2*. The study also showed that the CA-projecting Hug neurons may have a synaptic connection with the upstream Dh44 neurons.

## Materials and methods

### *Drosophila* strains

Flies were reared on 0.275 g agar, 5.0 g glucose, 4.5 g cornmeal, 2.0 g yeast extract, 150 μl propionic acid, and 175 μl 70% butylparaben (in EtOH) in 50 ml water. All experiments, except for the temperature shift experiment using *UAS-shibire*^*ts*^, were conducted at 25 °C under a 12:12-h light/dark cycle. For the temperature shift experiment, *UAS-shibire*^*ts*^ flies were first reared at 21 °C and transferred to 29 °C within 12 h after eclosion.

The following transgenic *D. melanogaster* strains were used in the study: *hug-YFP* (Melcher and Pankratz, 2005), *R18A04-GAL4* (BDSC #48793), *R65C11-LexA* (BDSC #54089), *R19D04-GAL4* (BDSC #48847), *hugS3-GAL4* (Melcher and Pankratz, 2005), *Dh44-R2-T2A-GAL4* (Kondo et al., 2020), *PK2-R2-T2A-GAL4* (Kondo et al., 2020), *PK2-R1-2A-GAL4* (BDSC # 84686), *LexAop-CD4::spGFP11,UAS-CD4::spGFP1-10* (BDSC #58755), *LexAop-mCherry* (BDSC #52272), *UAS-DenMark,UAS-Syt1::GFP; In(3L)D,mirr,D / TM6C* (BDSC #33064), *UAS-GFP;UAS-mCD8::GFP* (Ito et al., 1998) (a gift from Kei Ito), *UAS-TurboRFP* (a gift from A. Koto and M. Miura, The University of Tokyo), and *UAS-shibire*^*ts*^ (Kitamoto, 2001) (a gift from Hiroshi Kohsaka, The University of Tokyo).

### Generation of guinea pig anti-*Drosophila melanogaster* JH acid *O*-methyltransferase (JHAMT) antibody

Guinea pig polyclonal anti-*D. melanogaster* JHAMT antibody was raised against the peptide NH_2_-MNQASLYQHANQVQRHDAK-COOH, which corresponds to 1–19 amino acid residues of the mature JHAMT protein (Niwa et al., 2008).

### Immunohistochemistry

Tissues were dissected in phosphate buffer saline (PBS) and fixed in 3.7% formaldehyde in PBS for 30 to 60 min at room temperature (RT). The fixed samples were washed thrice in PBS, then washed for 15 min with PBS containing 0.3% Triton X-100 (PBT), and finally treated with the blocking solution (0.2% bovine serum albumin (Sigma A9647) in PBT) for 1 h at RT or overnight at 4 °C. Thereafter, the samples were incubated with a primary antibody in the blocking solution overnight at 4 °C. The primary antibodies used were as follows: mouse anti-GFP antibody (Sigma G6539, 1:2000); rabbit anti-RFP antibody (Medical & Biological Laboratories PM005, 1:2,000); rabbit anti-hugin-PK2 antibody (Schoofs et al., 2014) (1:500); rabbit anti-hugin-γ antibody (Schoofs et al., 2014) (1:500); and guinea pig anti-JHAMT antibody (this study, 1:1000-2000). After washing, the samples were incubated with the fluorophore (Alexa Fluor 488, 546, or 633)-conjugated secondary antibodies (Thermo Fisher Scientific) (1:200) in the blocking solution for 2 h at RT or overnight at 4 °C. After another round of washing, all samples were mounted on the glass slides using FluorSave reagent (Merck Millipore #345789).

### Quantitative RT-PCR for *Krüppel-homolog 1* (*Kr-h1*) expression

Total RNA was extracted from whole bodies of 4-day-old adult virgin female flies. RNA was reverse transcribed using ReverTra Ace qPCR RT Master Mix with gDNA remover (TOYOBO) and cDNA samples were used as templates for quantitative PCR using THUNDERBIRD SYBR qPCR mix (TOYOBO) on a Thermal Cycler Dice Real Time System (Takara Bio Inc.). The amount of target RNA was normalized to an endogenous control ribosomal protein 49 gene (*rp49*), and the relative fold change was calculated. The expression level of *Kr-h1* gene was compared using the ΔΔCt method. The following primers were used for *Kr-h1*, as previously described (Kang et al., 2017): *kr-h1 F* (5□-TCACACATCAAGAAGCCAACT-3□) and *kr-h1 R* (5□-GCTGGTTGGCGGAATAGTAA-3L). The primers for *rp49* were *rp49 F* (5□-CGGATCGATATGCTAAGCTGT-3□) and *rp49 R* (5□-GCGCTTGTTCGATCCGTA-3□).

### Statistical analysis

Sample sizes were chosen based on the number of independent experiments required for statistical significance and technical feasibility. The experiments were not randomized, and investigators were not blinded. All statistical analyses were performed using the “R” software environment.

## Results

### Some of the *Dh44-R2*-expressing neurons project to the corpora allata

This study originally intended to identify candidate neurons projecting to the ring gland (RG), comprised of the CA, CC, and PG, in *D. melanogaster* larvae, but not in the adult. We first browsed the large collections of larval images in the FlyLight database of Janelia Research Campus, Howard Hugh Medical Institute (https://flweb.janelia.org/cgi-bin/flew.cgi). The FlyLight database provides large anatomical image data sets and highly characterized collections of GAL4 lines that allowed us to visualize individual neurons in the *D. melanogaster* central nervous system (Jenett et al., 2012). In the FlyLight images of each GAL4 line, fluorescent proteins driven by the GAL4-UAS system (Brand and Perrimon, 1993) specifically visualize a subset of larval neurons, whereas many images do not include the RG located on the dorsal side of the two brain hemispheres. Nevertheless, some of the FlyLight images show neurons that extend their axons toward the longitudinal fissure of the two brain hemispheres, which are reminiscent of the axonal morphology of the identified RG-projecting neurons, such as PTTH neurons (McBrayer et al., 2007; Siegmund and Korge, 2001). Therefore, we expected that neurons that possess such characteristic axonal morphology are the RG-projecting neurons. Based on this working hypothesis, we could successfully identify a GAL4 line labeling a subset of the Crz neurons that projects to the prothoracic gland cells (Imura et al., 2020).

During the course of our FlyLight database search, we found not only the *Crz-GAL4* line projecting to the prothoracic gland but also other GAL4 lines that visualize neuronal processes at the longitudinal fissure of the two brain hemispheres (E.I, Y.M, Y.K., Y.S.N., R.N., unpublished). Next, we obtained the GAL4 lines, and crossed them with *UAS-mCD8::GFP* strain to examine whether the GFP-labeled neurons project to the RGs in the 3^rd^-instar larvae. These strains included *R18A04-GAL4*, in which *GAL4* expression was driven by a part of the enhancer region of the G protein-coupled receptor gene *Dh44-R2* (Figure 1a). We confirmed that *R18A04-GAL4* was active in neurons projecting to the central region of the RG, which seemed to correspond to the CA (Figure 1b). To precisely examine whether the *R18A04-GAL4*-positive neurons projected to the CA, by immunostaining, we generated a new guinea pig antibody against JHAMT protein (Niwa et al., 2008). We found GFP signals in *R18A04-GAL4> mCD8::GFP* larvae to be observable in the neurite terminals located at the CA, which was JHAMT-positive (Figure 1b).

**Figure 1.**
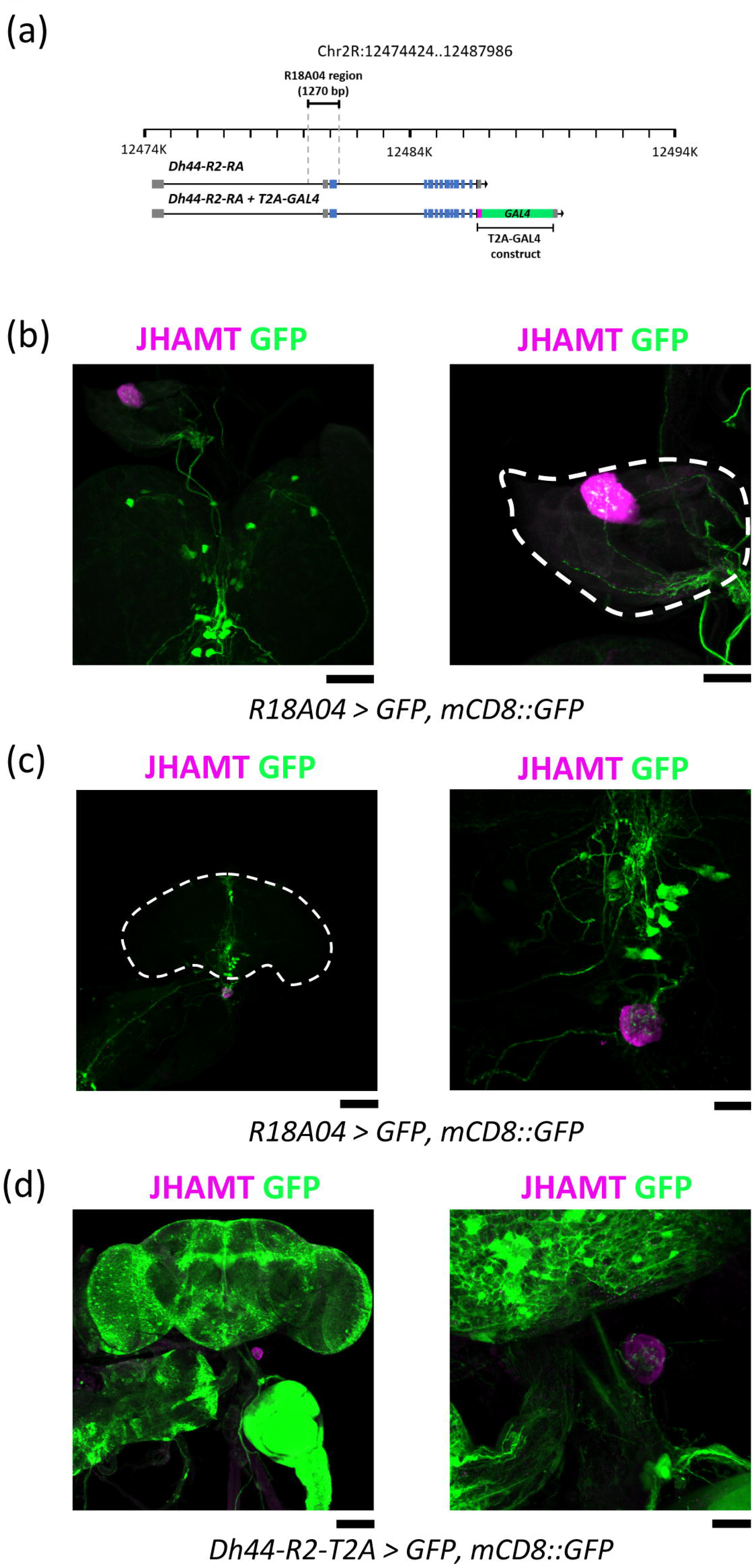
*Dh44-R2*-expressing neurons project to the CA. (a) The enhancer region of *R18A04-GAL4* and the construct of *Dh44-R2-T2A-GAL4*. Gray, blue, magenta, and green boxes indicate the untranslated region, coding sequence, T2A peptide sequence, and *GAL4* sequence, respectively. (b) The brain-RG complex (left) and the RG (right) of *R18A04-GAL4; UAS-*GFP, *UAS-mCD8::GFP* in the 3^rd^-instar larval stage. Scale bar, 50 μm (left), 25 μm (right). (c) *R18A04-GAL4*-positive neurons project to the CA in the adult stage. The brain-proventriculus (PV) complex (left) and the CA (right). Scale bar, 100 μm (left), 25 μm (right). (d) *Dh44-R2-T2A-GAL4*-positive neurons project to the CA. The brain-PV complex (left) and the CA (right). Scale bar, 100 μm (left), 25 μm (right).

Notably, we found the projection of *R18A04-GAL4*-positive neurons to the CA to be observable in the adult stage (Figure 1c). Till date, CA-projecting neurons had not been identified and characterized in adult *D. melanogaster*. On the other hand, many recent studies have reported JH biosynthesis to be adaptively regulated in response to internal physiological conditions and external environmental cues in adult *D. melanogaster* (Lee et al., 2017; Meiselman et al., 2017; Ojima et al., 2018; Reiff et al., 2015; Wu et al., 2018; Yamamoto et al., 2013). Therefore, we decided to focus on the anatomical characteristics of *R18A04-GAL4*-positive neurons in the adult stage.

Since the expression of *R18A04-GAL4* is regulated by the enhancer region of *Dh44-R2*, we expected the *R18A04-GAL4*-positive CA-projecting neurons to express *Dh44-R2* gene. To test this idea, we used the *Dh44-R2* knock-in T2A-GAL4 line (Kondo et al., 2020), and found a group of neurite termini, labeled with GFP of *Dh44-R2-T2A-GAL4> UAS-mCD8::GFP*, to be located at the CA in the adult stage (Figure 1d). These results clearly suggested that neurons expressing *Dh44-R2* project to the CA.

Since Dh44-R2 is a receptor for the neuropeptide Dh44, CA-projecting *Dh44-R2*-expressing neurons might receive a neuronal input from Dh44 neurons. To confirm the anatomical relationship of the ligand-expressing neurons with receptor-expressing neurons, we performed co-labeling experiment using *R65C11–LexA* and *R18A04-GAL4*. Since *R65C11–LexA* is a fly strain that expresses *LexA* transgene under control of *Dh44* promoter, hereafter, we refer to *R65C11–LexA* as *Dh44-LexA*. We found both *Dh44-LexA*-labeled and *R18A04-GAL4*-labeled neurites to be in close proximity in the SEZ of the adult brain (Figure 2a). The GFP Reconstitution Across Synaptic Partners (GRASP) method (Feinberg et al., 2008) was used to confirm whether the neurons were close enough in proximity to form synapses. As expected, we observed reconstituted GFP fluorescence at the sites where neurites were observed (Figure 2a,b). These results suggested that the CA-projecting *Dh44-R2*-expressing neurons have direct synaptic contact with the upstream Dh44 neurons.

**Figure 2.**
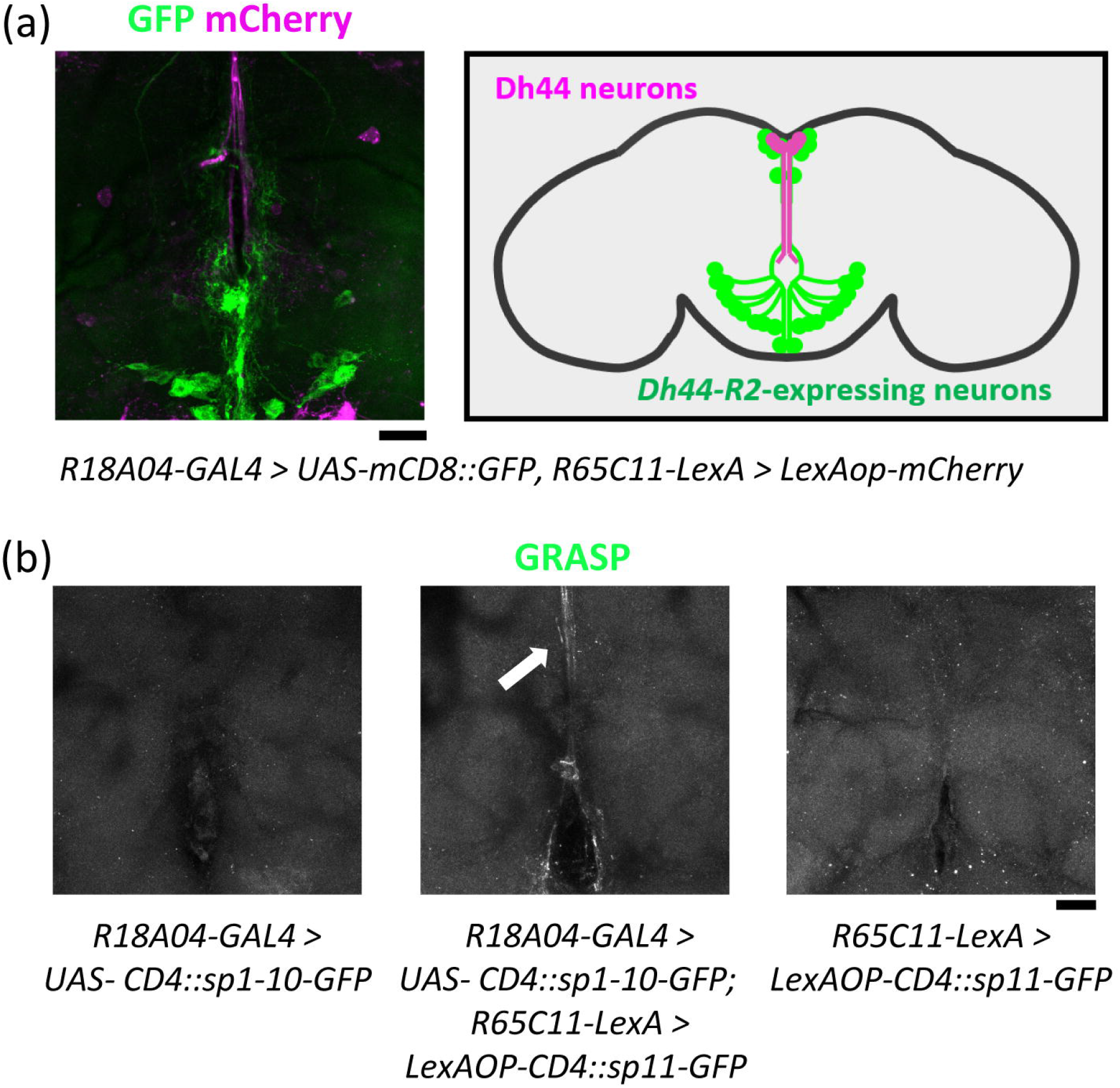
Dh44 neurons and *Dh44-R2* neurons have synaptic contact. (a) *Dh44-R2* neurons are adjacent to Dh44 neurons. Left: Co-staining of Dh44 neurons (magenta) and *Dh44-R2* neurons (green). Scale bar, 25 μm. Right: Schematic diagram showing the cell bodies and neurites of Dh44 neurons (magenta) and *Dh44-R2* neurons (green). (b) In GRASP analysis, *spGFP1-10::CD4* was expressed by *R18A04-GAL4*, and *spGFP11::CD4* was expressed by *R65C11–LexA* (middle panel). GRASP signal was detected between *Dh44-R2* neurons and Dh44 neurons (arrow). In contrast, negative controls did not give signals, as shown in left and right panels. Scale bar, 25 μm.

### The corpora allata-projecting *Dh44-R2*-expressing neurons produce the neuropeptide hugin

We next investigated which neurotransmitter is utilized in the CA-projecting *Dh44-R2*-expressing neurons. For this analysis, we focused on a subset of neurons producing the neuropeptide Hug, which is synthesized in approximately 20 neurons, whose cell bodies are located in the SEZ of the *D. melanogaster* brain (Melcher and Pankratz, 2005; Meng et al., 2002). Previous studies had reported that there are several clusters of Hug neurons, each of which projects to various regions, including not only the pars intercerebralis and ventral nerve cord but also the CC-CA complex in the adults (King et al., 2017; Melcher and Pankratz, 2005). However, the previous study had not clarified which endocrine organs the Hug neurons project to, the CA or CC. Through our anatomical analysis of *R18A04-GAL4* and *Dh44-R2-T2A-GAL4* flies, we hypothesized that overall morphology of the CA-projecting *Dh44-R2*-expressing neurons is similar to that of the CA-projecting Hug neurons. To test this hypothesis, we simultaneously labeled both *Dh44-R2*-expressing neurons and Hug neurons with *R18A04-GAL4*-or *Dh44-R2-T2A-GAL4-*driven red fluorescence protein (RFP) and *hug* promoter-driven yellow fluorescence protein (YFP), respectively. We found that, with either of *R18A04-GAL4* or *Dh44-R2-T2A-GAL4*, GAL4-driven RFP signal was observed in the cell bodies of *hug*-promoter-driven YFP signal (Figure 3a,c). Moreover, we observed both YFP and RFP signals in the identical axon terminals located on the CA (Figure 3b,d). Together, the results suggest that the CA-projecting *Dh44-R2*-expressing neurons are indeed Hug-positive.

**Figure 3.**
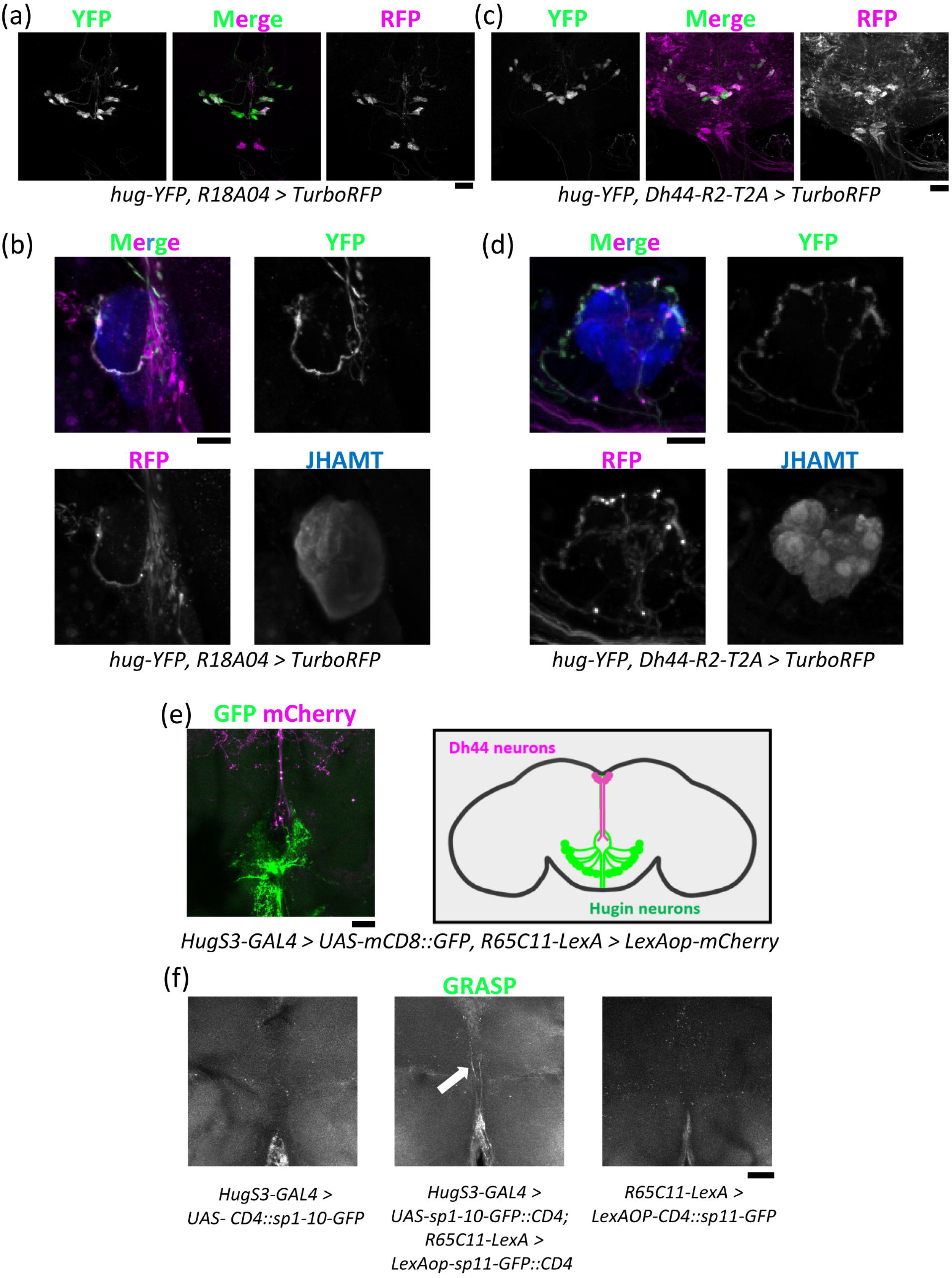
The CA-projecting *Dh44-R2*-expressing neurons produce Hug. (a) *R18A04-GAL4; UAS-TurboRFP* signals (magenta) are detected in the cell bodies of Hug neurons (green). Scale bar, 25 μm. (b) RFP signals reflecting *R18A04-GAL4* expression are merged with Hug-YFP signals (green) in the axon terminals. Scale bar, 10 μm. (c) *Dh44-R2-T2A-GAL; UAS-TurboRFP* signals (magenta) are detected in the cell bodies of Hug neurons (green). Scale bar, 25 μm. (d) RFP signals reflecting *Dh44-R2* expression are merged with Hug-YFP signals (green) in the axon terminals. Scale bar, 10 μm. (e) Hug neurons are adjacent to Dh44 neurons. Left: Co-staining of Dh44 neurons (magenta) and Hug neurons (green). Scale bar, 25 μm. Right: Schematic diagram showing the cell bodies and neurites of Dh44 neurons (magenta) and Hug neurons (green). (f) In GRASP analysis, *spGFP1-10::CD4* was expressed by *hugS3-GAL4*, and *spGFP11::CD4* was expressed by *R65C11–LexA* (middle panel). GRASP signal was detected between Hug neurons and Dh44 neurons (green, arrow). In contrast, negative controls did not give signals, as shown in left and right panels. Scale bar, 25 μm.

To confirm whether the Dh44 neurons are located upstream of the Hug neurons, we performed the same experiments as in Fig. 2 using *Dh44-LexA* along with *hugS3-GAL4* strain, in which *GAL4* is expressed downstream of the *hug* promoter (Melcher and Pankratz, 2005). We found the neurites of Hug neurons labeled with *hugS3-GAL4* to be closely located to those of the Dh44 neurons labeled with *Dh44-LexA* (Figure 3e). Furthermore, GRASP experiments showed a reconstituted GFP fluorescence at the sites where both Hug and Dh44 neurites were observed (Figure 3f). The results, therefore, suggest that the CA-projecting Hug neurons have direct synaptic contact with the upstream Dh44 neurons.

### The corpora allata-projecting hugin neurons may be dispensable for juvenile hormone biosynthesis

As shown above, *R18A04-GAL4* line was active in at least 14 Hug neurons in the SEZ (Figure 3a). Given the fact that Hug neurons divide into multiple subpopulations that project to distinct regions (Schlegel et al., 2016), we expected a few, not all, of these Hug neurons to project to the CA. This expectation was supported by our anatomical observation using another GAL4 line that we identified during our FlyLight image screen, namely *R37F05-GAL4*, which is under the control of a small enhancer region of retinal degeneration A gene. We found that, while *R37F05-GAL4* was active in several neurons, only 2 pairs of the neurons were merged with *hug* promoter-driven fluorescence signal (Figure 4a). Importantly, morphology of the *R37F05-GAL4* neurite termini on the CA was identical to that of the CA-projecting Hug neurons (Figure 4b). Moreover, *R37F05-GAL4*-driven synaptotagmin-GFP (Syt1::GFP), a synaptic vesicle marker, was observed on the CA while *R37F05-GAL4*-driven dendritic marker DenMark was not (Figure 4c). These results strongly suggest that only the 2 pairs of Hug neurons, not the rest, project to the CA in adults.

**Figure 4.**
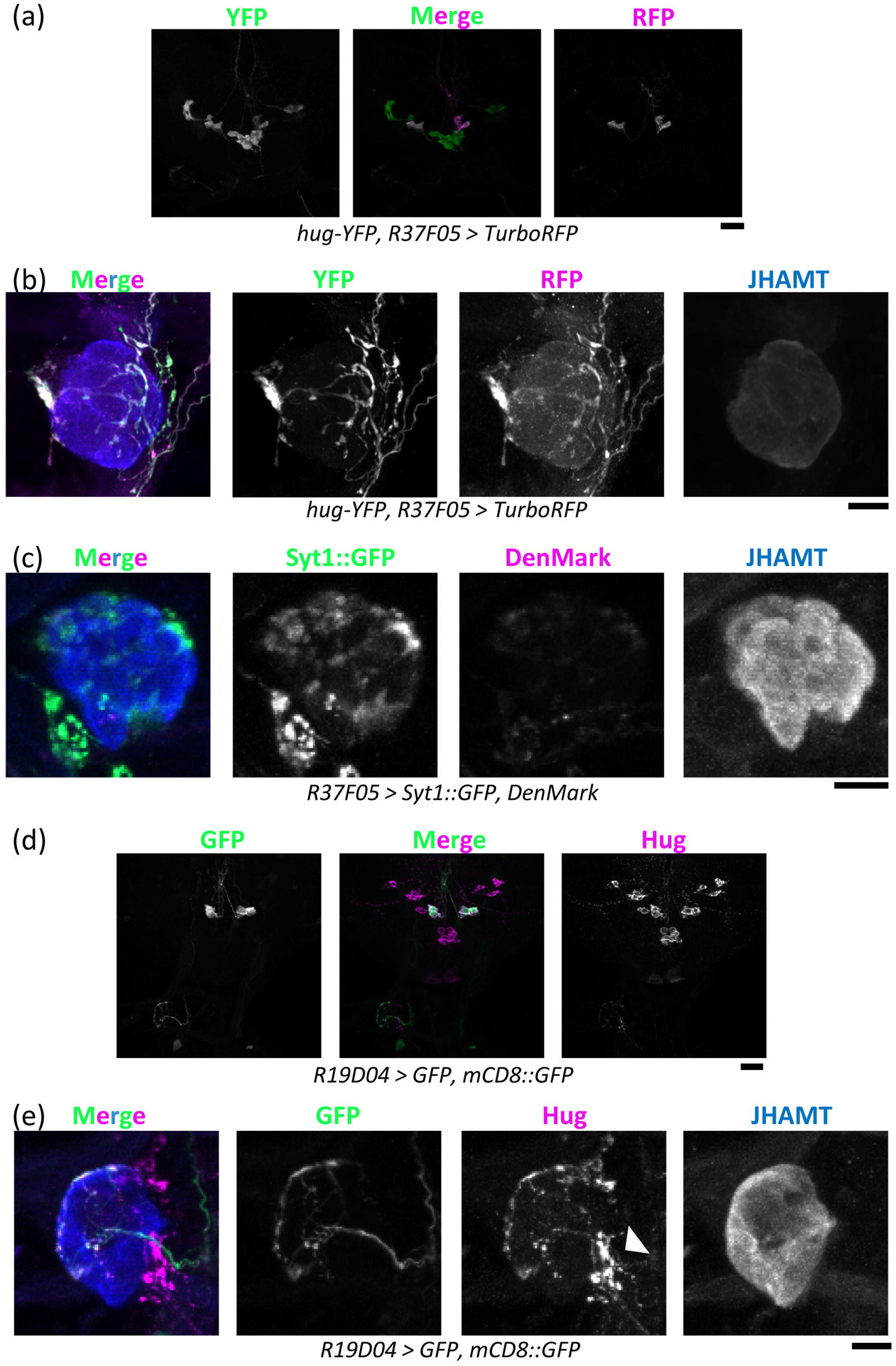
Two pairs of Hug neurons project to the CA. (a) *R37F05-GAL4; UAS-TurboRFP* signals (magenta) are detected in the cell bodies of Hug neurons (green). Scale bar, 25 μm. (b) RFP signals reflecting *R37F05-GAL4* expression are merged with Hug-YFP signals (green) in the axon terminals. Scale bar,10 μm. (c) A synaptic vesicle marker Syt1::GFP, but not a dendritic marker Denmark, was observed on the CA. *Syt1::GFP* and *DenMark* were expressed by *R37F05-GAL4*. Scale bar, 25 μm. (d) Cell bodies of the CA-projecting Hug neurons are overlapped with those of *R19D04-GAL4*-positive neurons. Scale bar, 25 μm. (e) *R19D04-GAL4; UAS-GFP*, and *UAS-mCD8::GFP* neurons (green) are merged with Hug neurons (magenta). Anti-Hug antibody labeled not only the CA-projecting neurons but also neurons projecting to regions other than the CA (arrowhead). Scale bar, 10 μm.

Further, we realized by chance that another GAL4 line, namely *R19D04-GAL4*, which is under the control of a small enhancer region of *asense* gene, also labeled the 2 pairs of CA-projecting Hug neurons in the adult (Figure 4d) while we did not focus on *R19D04-GAL4* line at our initial FlyLight image screen. We confirmed that *R19D04-GAL4*-positive neurons indeed project to the CA (Figure 4e).

Since *R19D04-GAL4* is active in fewer neurons than other GAL4 lines used in this study, we utilized *R19D04-GAL4* to manipulate neuronal activity of the CA-projecting Hug neurons in the following experiment. We expressed the gene encoding temperature-sensitive dynamin (*shibire*^*ts*^) specifically in the CA-projecting Hug neurons and inhibited their neuronal activity specifically in the adult stage at a restrictive temperature (Kitamoto, 2001). We collected mRNA from these flies and examined expression of the *Kr-h1* gene; *Kr-h1* is well known as a downstream target of JH receptors and its expression level correlates well with hemolymph JH titer level (Minakuchi et al., 2009). However, we found the inhibition of CA-projecting Hug neurons to not significantly alter *Kr-h1* expression compared to that in the controls (Figure 5a).

**Figure 5.**
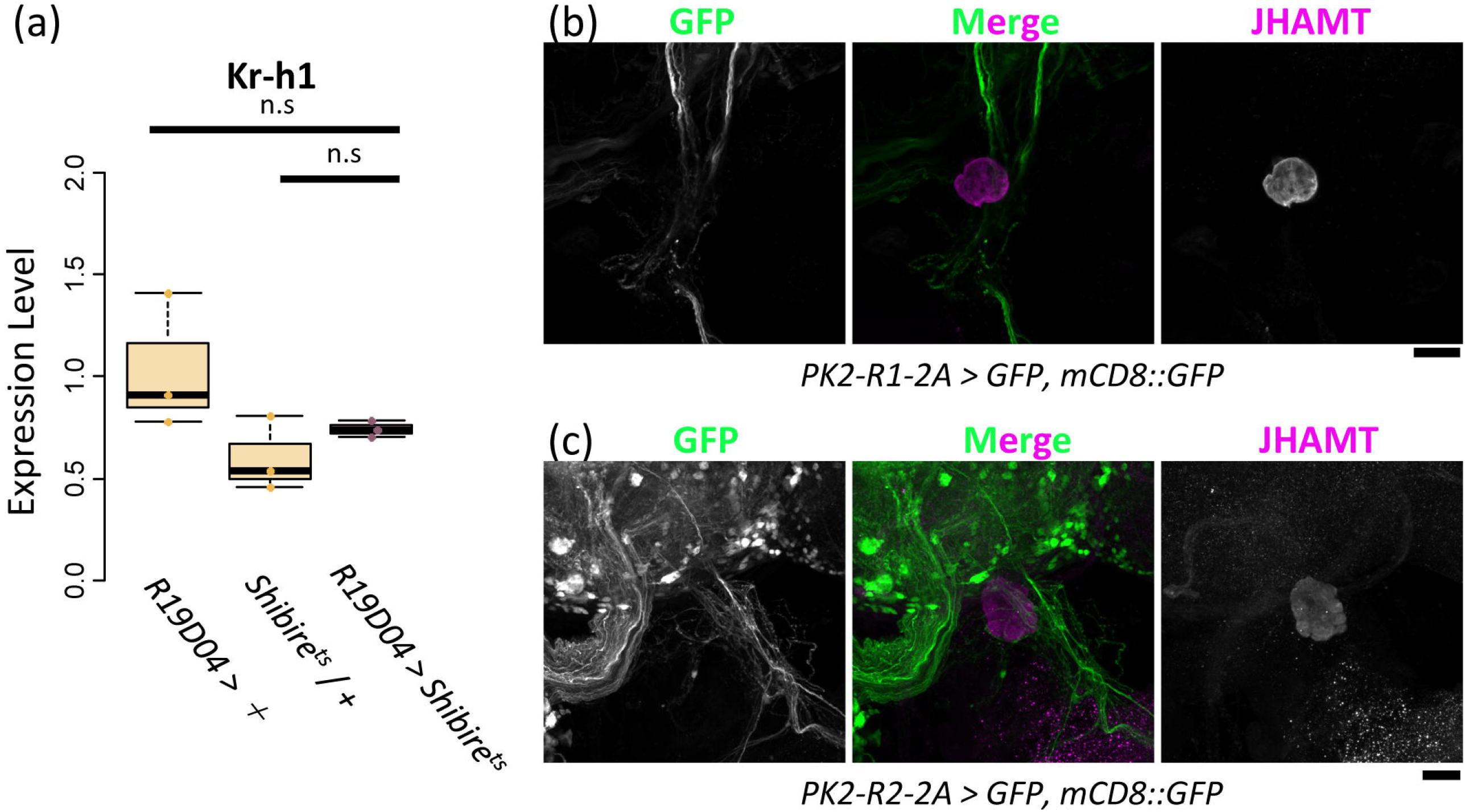
The CA-projecting Hug neurons might not be required for JH biosynthesis. (a) The *shibire*^*ts*^-mediated neuronal inhibition of CA-projecting Hug neurons could not change the expression level of *Kr-h1*. Student’s *t*-test with Bonferoni’s collection was used for this data. \*\*\**p* ≤ 0.001, \*\**p* ≤ 0.01, and \**p* ≤ 0.05; n.s., non-significant (*p* > 0.05). (b, c) GFP signals reflecting *PK2-R1* (b) and *PK2-R2* (c) expression are not detected in the CA. Scale bar, 25 μm.

We also took another approach to examine whether Hug signaling is involved in the regulation of JH biosynthesis in the CA. A previous study had identified 2 specific receptors for Hug, namely Pyrokinin 2 Receptor 1 (PK2-R1) and Pyrokinin 2 Receptor 2 (PK2-R2) (Park et al., 2002). To examine whether these receptors are expressed in the CAs, we performed cell-specific labeling using *PK2-R1-T2A-GAL4* and *PK2-R2-T2A-GAL4* lines. However, we could not observe any significant *T2A-GAL4*-driven GFP fluorescence signal in the CA (Figure 5b,c). In conjunction with our observation regarding *Kr-h1* expression (Figure 5a), these results suggested the CA-projecting Hug neurons to be dispensable for JH biosynthesis; they may rather play a role in systemic function of *D. melanogaster*, which will be discussed later.

## Discussion

In this study, we anatomically characterized the CA-projecting Hug neurons in *D. melanogaster*. Hug was originally identified as a myostimulatory and ecdysis-modifying neuropeptide (Meng et al., 2002); further studies have demonstrated that it plays a central role in integrating external and internal feeding-relevant information to control feeding behavior (Bader et al., 2007; Hückesfeld et al., 2016; Melcher and Pankratz, 2005; Melcher et al., 2006; Melcher et al., 2007; Schoofs et al., 2014; Surendran et al., 2017). More recently, Hug has been found to regulate circadian rhythm in *D. melanogaster* (King et al., 2017). To understand the role of Hug in *D. melanogaster* physiology and behavior, neuronal circuits and synaptic connections of Hug neurons have been intensively studied (Bader et al., 2007; Melcher and Pankratz, 2005; Schlegel et al., 2016). Regarding the anatomical relationship between Hug neurons and the CA, the former have been reported to project to the complex including the CA and CC in the adult stage (Melcher and Pankratz, 2005), and to the CC in the larval stage (Hückesfeld et al., 2020). However, these studies have not precisely examined whether a part of Hug neurons project to the CA. To the best of our knowledge, this is the first study to identify a specific neurotransmitter that is produced in the CA-projecting neurons of *D. melanogaster*, and to anatomically characterize a neuronal relay to the CA-projecting neurons from the upstream interneurons.

Based on our anatomical observation, we expected the CA-projecting Hug neurons to be involved in the regulation of JH biosynthesis in the CA. However, our genetic analysis till date has not validated this expectation. Particularly, to our surprise, either of the 2 identified Hug receptor genes, *PK2-R1* and *PK2-R2*, appeared not to be expressed in the CA. Although it has been reported that another receptor called PK1-R is potentially activated by Hug in a heterologous cell culture experiment (Cazzamali et al., 2005), the half-maximal effective concentration of hug for PK1-R is more than 10 μM (Park et al., 2002). Therefore, we have not examined the involvement of PK1-R in this study. Alternatively, an unknown Hug receptor may still possibly exist and its gene expressed in the CA. Nevertheless, our data implied that Hug released from the CA-projecting Hug neurons is not locally received by the CA cells, and rather received by other tissues apart from the CA. In this scenario, the CA-projecting Hug neurons would secrete Hug from the synaptic termini on the CA to hemolymph. Such systemic release of neuropeptides from the CA is already known in the silkworm *Bombyx mori*, since *B. mori* PTTH neurons project to the CA instead of the PG, and the secreted PTTH from the CA is circulated in the hemolymph and then received by the PG cells (Mizoguchi et al., 1990). Whether the CA-projecting neuron-derived Hug is released to hemolymph and, if so, how essential the secreted Hug is for regulating *D. melanogaster* biological functions, other than JH biosynthesis, would be worth investigating.

Our current data suggest that the CA-projecting Hug neurons have synaptic connections with the upstream Dh44 neurons. Dh44 is a crucial neuropeptide that regulates the rest:activity rhythms and sleep in *D. melanogaster* (Cavanaugh et al., 2014). In addition, Dh44 receptor 1 (*Dh44-R1*), though not *Dh44-R2*, is involved in the regulation of circadian locomotor activity and is expressed in Hug neurons in the SEZ of *D. melanogaster* (King et al., 2017). Therefore, it is feasible to think that, analogous to the neuronal relay from the Dh44 neurons to *Dh44-R1*-expressing neurons, *Dh44-R2*-expressing neurons might be involved in the regulation of circadian rhythms and/or sleep. Of note, JH is involved in the regulation of sleep behavior in *D. melanogaster* (Wu et al., 2018) and of circadian rhythms in the bumble bee *Bombus terrestris* (Pandey et al., 2020). Alternatively, since Dh44 neurons are also important for the sensing mechanisms of dietary sugar and amino acids (Dus et al., 2015; Yang et al., 2018), the CA-projecting Hug neurons might play a role in nutritional sensing and metabolism. JH biosynthesis has also been suggested to be dependent on the nutritional status of some insects (Badisco et al., 2013).

We should also note that the CA-projecting Hug neurons that we identified might be different from the previously identified CA-projecting neurons, namely CA-LP1 and CA-LP2, in *D. melanogaster* (Siegmund and Korge, 2001). While the cell bodies of the CA-projecting Hug neurons are located in the SEZ, those of CA-LP1 and CA-LP2 are placed in the lateral protocerebrum (Siegmund and Korge, 2001). Although the role of CA-LP1 and/or CA-LP2 might suppress JH biosynthesis in the CA to regulate epithelial morphogenesis of male genitalia during pupal stage (Ádám et al., 2003), which neurotransmitters are produced in CA-LP1 and CA-LP2 have not been elucidated yet. In future, identities of these CA-projecting neurons should be analyzed, and the functional relationship across CA-LP1, CA-LP2, and Hug neurons should be investigated to understand the molecular and neuronal mechanisms of JH biosynthesis.

## Acknowledgements

We thank Kei Ito, Hiroshi Kohsaka, Akiko Koto, Masayuki Miura, the Bloomington Stock Center, the Kyoto Stock Center (DGRC), the National Institute of Genetics, the Vienna Drosophila Resource Center, and the Developmental Studies Hybridoma Bank for providing all stocks and reagents. We would like to thank Editage (www.editage.com) for English language editing. E.I. was a recipient of fellowship from the Japan Society for the Promotion of Science. This work was supported by grants from PRESTO of Japan Science and Technology Agency to R.N., from Japan Society for the Promotion of Science KAKENHI 17J00218 to E.I., KAKENHI 26250001 to H.T., and A17H01378 to H.T., and from the Tomizawa Jun-ichi & Keiko Fund of Molecular Biology Society of Japan for Young Scientist to Y.S.N.

